# A novel nested gene *Aff3ir* participates in vascular remodelling by enhancing endothelial cell differentiation in mice

**DOI:** 10.1101/2023.12.09.570916

**Authors:** Yue Zhao, Mazdak Ehteramyan, Yi Li, Xuefeng Bai, Lei Huang, Yingtang Gao, Angshumonik Angbohang, Xiaoping Yang, Steven Lynham, Andriana Margariti, Ajay M Shah, Yaling Tao, Ting Cai, Tong Li, Min Zhang, Lingfang Zeng

## Abstract

Endothelial integrity in the vasculature is critically maintained by vascular stem/progenitor cells (SPCs) giving rise to endothelial cells (ECs). However, the genes significantly activated during differentiation remain incompletely understood. Based on mouse aorta and vein cDNA library, we unearthed a hitherto unidentified gene nested residing within intron 6 of *Aff3*, christened as *Aff3* intron resident (*Aff3ir*), upregulated during laminar shear stress-induced ECs differentiation in mouse. Proteomic analysis substantiated the presence of a 45-amino acid(aa) peptide (AFF3IR-ORF1) and 109-aa or 151-aa protein (AFF3IR-ORF2) encoded from two transcript variants. During embryonic development, AFF3IR-ORF1 peaked at E14.5, while AFF3IR-ORF2 displayed a continuous increase until E19.5. In adult mice, AFF3IR-ORF1 was detected in the lung, liver, spleen, and kidney, while AFF3IR-ORF2 was most abundant in the aorta. Furthermore, Western blot and immunofluorescence analyses revealed a specific upregulation of AFF3IR-ORF2, but not AFF3IR-ORF1, three days after femoral artery injury or hindlimb ischemia *in vivo*. Overexpression of AFF3IR-ORF2 enhanced, while its knockdown attenuated, SPCs differentiation into ECs induced by shear stress or vascular endothelial growth factor *in vitro*. Notably, the upregulated AFF3IR-ORF2 hindered SPCs proliferation by sequestering minichromosome maintenance complex component 3 in the cytoplasm, thereby shifting the status of SPCs from a pro-proliferation to a pro-differentiation state. In conclusion, our discoveries unveil the novel protein-coding gene *Aff3ir* as a participant in ECs differentiation, providing fresh insights into the regulation of vascular endothelial integrity.

## Introduction

Endothelial integrity stands as a pivotal determinant in upholding cardiovascular homeostasis. In the aftermath of endothelial impairment or denudation resulting from vascular injury, the restoration of endothelium integrity emerges as a paramount stage. This intricate process predominantly encompasses the proliferation and migration of neighbouring undamaged endothelial cells (ECs), accompanied by the orchestration of resident and bone marrow-derived vascular stem/progenitor cells (SPCs), characterized by activation, mobilization, and subsequent differentiation^1^. Over the course of four decades, the evolution of regenerative medicine within the realm of SPCs has steadily gained prominence as a promising therapeutic avenue aimed at addressing a spectrum of vascular afflictions, spanning coronary artery disease, stroke, and peripheral arterial disorders.

In response to dynamic cues like shear stress^2^ or signalling molecules such as vascular endothelial growth factor (VEGF)^3^, a repertoire of genes becomes activated, orchestrating the transformative journey of SPCs into mature ECs. A subset of these genes has been duly recognised for their pivotal roles in reendothelialization^4–6^, emblematic of the restoration of the endothelial layer following vascular injury. Our previous studies and other teams have also discerned plenty of crucial contributors, including histone deacetylases^7–9^ and SOX18^10^, which have provided insights into the complex molecular framework supporting this transformative process. Nevertheless, a more intricate tapestry emerges when scrutinizing gene screenings, revealing an extensive array of genes experiencing dynamic regulation during the differentiation of stem cells^11, 12^. The genomic landscape alludes to a nuanced interplay that encompasses a multitude of genes, thereby compelling a more comprehensive inquiry into the molecular signatures that govern this process.

In a prior study, we unveiled the heightened expression of a distinctive cDNA fragment sourced from the comprehensive mouse aorta and vein cDNA library. This elevation coincided with the progressive differentiation of SPCs into ECs under the influence of laminar flow stress in the murine context. Building upon this discovery, we present here a novel nested gene christened as *Aff3ir*. This hitherto unexplored genetic entity emerges as a pivotal player in the realm of vascular dynamics, fostering the perpetuation of endothelial integrity. Central to its impactful role is one of its encoded proteins, named as AFF3IR-ORF2, which emerges as a protagonist in enhancing the process of EC differentiation in the murine model. This finding unfolds a fresh vista of molecular intricacies, hinting at the potential of *Aff3ir* and its encoded protein AFF3IR-ORF2 to substantially contribute to the sustenance of vascular homeostasis.

## Results

### The discovery of a novel nested gene Aff3ir

In our previous investigation, we demonstrated that laminar shear stress serves as a driving force for EC differentiation, originating from mouse embryonic stem cells (mESCs) that have undergone 3-days spontaneous differentiation into SPCs^9^. To delve deeper into the mechanistic aspects, we conducted a microarray analysis encompassing gene expression profiles of murine ESCs and their spontaneously differentiated counterparts. These cell populations were subjected either to a laminar shear stress of 12 dynes/cm² or were maintained under static conditions. Notably, we pinpointed an intriguing 874bp cDNA fragment sourced from the mouse aorta and vein cDNA library^13^ (*GenBank: AK040668.1*), which exhibited a conspicuous upregulation under conditions of shear stress (Fig 1A). Upon subjecting this fragment to a comparative genomic examination, we discerned its presence within intron 6 of the mouse *Aff3* gene, encompassing distinct genomic domains.

**Figure 1.**
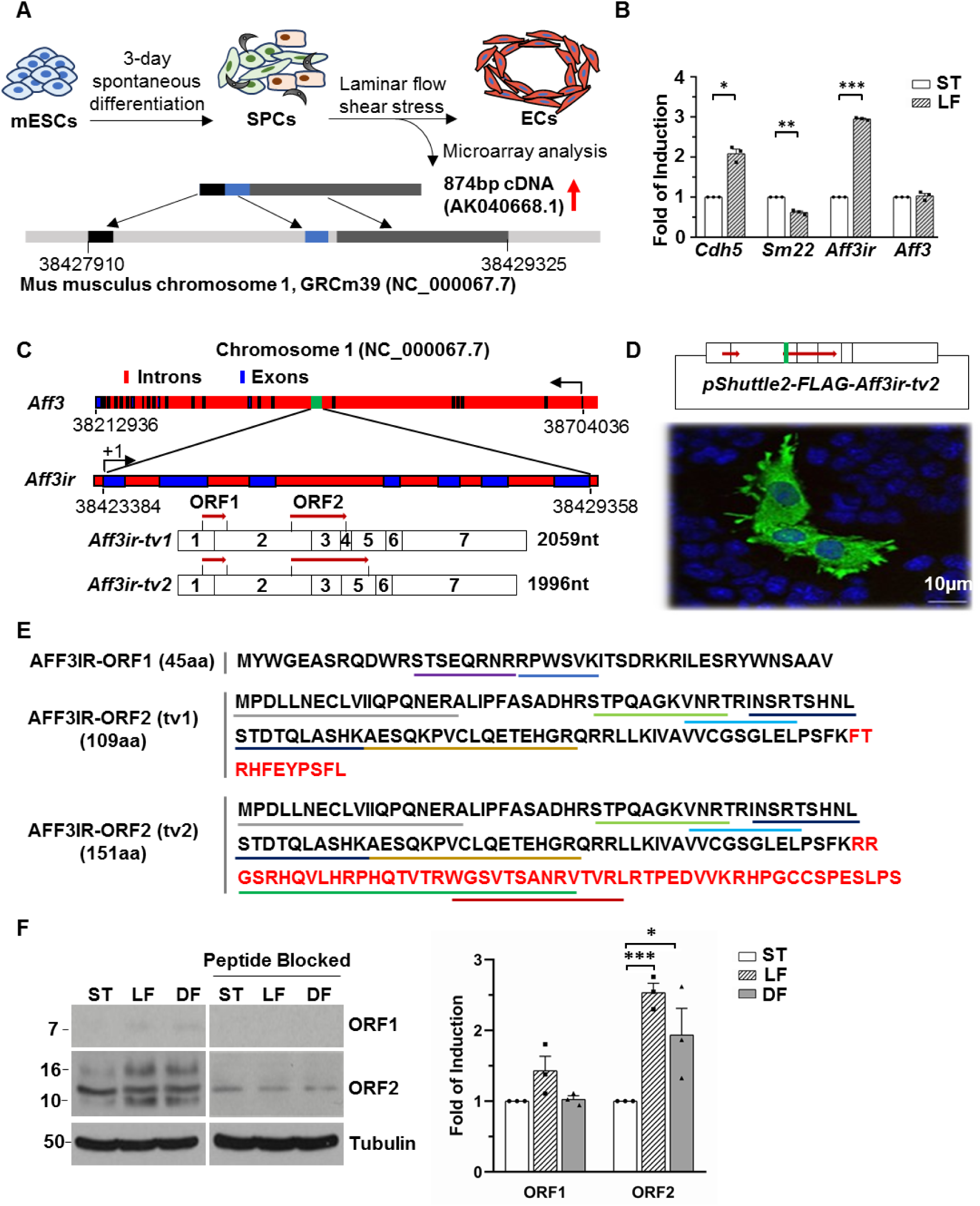
Discovery of the novel protein-coding gene *Aff3ir*. (**A**) A brief chart flow of the discovery process. (**B**) qRT-PCR analysis of the relative expression of the candidate genes in mouse stem/progenitor cells (SPCs) that were derived from 3-day spontaneously differentiated embryonic stem cells (ESCs) followed by 24hr laminar shear stress with static control included (n=3). (**C**) A schematic illustration of the *Aff3ir* gene structure. *Aff3ir* (green box) is in intron 6 of *Aff3*, comprising of 7 exons (blue box). Number shows the nucleotide position in chromosome 1. +1/arrow indicates the transcription initiation site and direction. Dark red arrows indicate the predicted short open reading frames (ORF). (**D**) SPCs were transfected with *pShuttle2-Aff3ir-FLAG-tv2* plasmid (upper, number in box shows the exons. Brown arrows show the ORFs The green box indicates the inserted FLAG in ORF2), followed by IF with anti-FLAG antibody (lower), DAPI included to counterstain nuclei. (**E**) Computational deduction of the amino acid sequences for AFF3IR-ORF1 and AFF3IR-ORF2 with the same N-terminal sequence indicated in black for proteins derived from *Aff3ir-tv1* and *Aff3ir-tv2*. Coloured underlines indicate peptide sequences detected by mass spectrometry. (**F**) The Sca1^+^-VPCs were subjected to laminar flow (LF), disturbed flow (DF) in the dish-shaking system or kept at static condition (ST) for 24hr, followed by WB analysis with anti-AFF3IR-ORFs or blocking peptide (left) and quantification (right). Data presented were representative images or mean SEM using one-way ANOVA with GraphPad Prism 8 multiple comparison test of three independent experiments. *: p<0.05, ***: p<0.001.

The investigation of homologous counterparts brought forth the identification of a parallel fragment located within intron 7 of the human *AFF3* gene. The homologous sequences were traced across diverse human tissues: *AI438968* sourced from B-cells of chronic lymphatic leukemia, *AA541378* sourced from metastatic prostate lesions of the bone, and *BF751024* sourced from normal breast tissue. Termed as *Aff3* intron resident gene or *Aff3ir*, this 874bp fragment emanates the possibility of being a derivative of the *Aff3* precursor RNA or possibly transcribed independently.

To scrutinize this scenario, we devised specific primers for *Aff3ir* and *Aff3*, followed by quantitative reverse transcription polymerase chain reaction (qRT-PCR) assessment. This experimental course was enacted using mESCs derived SPCs, which were alternately exposed to 12 dynes/cm² laminar shear stress or retained under static conditions. Strikingly, our findings unveiled a significant upregulation of *Aff3ir* while *Aff3* unchanged, in consonance with the EC marker *Cdh5*, exclusively in response to laminar flow (Fig. 1B). This observation posits that *Aff3ir* and *Aff3* exhibit separate transcriptional activities under the influence of shear stress, thereby suggesting a plausible link between *Aff3ir* and the process of EC differentiation.

To meticulously capture the terminal sequences at both the 3’ and 5’ ends, we employed the rapid amplification of cDNA ends technique. Subsequent DNA sequencing scrutiny unveiled two distinctive clones, each measuring 2059nt (Fig. S1A, Genbank: MH282850.1) and 1996nt (Fig. S1B, Genbank: MH282851.1) in size with the latter clone featured an internal omission of 63 nucleotides. A comparative alignment of these clone sequences against the expansive canvas of the mouse genome exposed the existence of divergent transcript variants, denominated as *Aff3ir-tv1* and *Aff3ir-tv2*, respectively. Notably, the structural arrangement of these variants diverges; *Aff3ir-tv1* spans 7 exons, whereas *Aff3ir-tv2* comprises 6 exons with the absence of exon 4 (Fig. 1C). The predominant presence of *Aff3ir-tv2* emerged evident, as it surpassed *Aff3ir-tv1* by a considerable margin, maintaining a ratio of 50:1. Further anatomization of the *Aff3ir* gene architecture unveiled that *Aff3ir* and *Aff3* engaged disparate DNA strands for their transcriptional activities, thereby underscoring their unique and divergent transcriptional profiles (Fig. 1C).

### Aff3ir is an encoding duo-cistronic gene

The predominant paradigm of protein-coding overlapping genes has been the nested gene configuration, residing entirely within an intron of an external gene, a phenomenon well-established previously^14^. In pursuit of understanding the potential encoding capacity of *Aff3ir*, online computational modelling tools (https://web.expasy.org/translate/) were embarked. We foretold the existence of multiple open reading frames (ORFs) within the *Aff3ir* locus (Fig. 1C). Specifically, ORF1 encompasses a 45-amino acid (aa) sequence across both transcript variants (*tv*), while ORF2 furnishes proteins spanning 109-aa from *Aff3ir-tv1* and 151-aa from *Aff3ir-tv2* (Fig. 1C). Noteworthy is the shared N-terminal region between the two proteins, diverging only in their C-terminal segments due to the presence of a stop codon within exon 4 of *Aff3ir-tv1*.

To unravel the encoding potential of *Aff3ir*, we initiated the generation of a *pShuttle2-FLAG-Aff3ir-tv2* plasmid. This construct seamlessly incorporated the entire *Aff3ir-tv2* cDNA sequence into the *pShuttle2* vector, with a strategically positioned FLAG sequence situated downstream of the AUG codon within ORF2 (Fig. 1D, upper segment). Subsequent immunofluorescence (IF) staining assays distinctly illuminated a robust signal for the FLAG epitope in SPCs transfected with *pShuttle2-FLAG-Aff3ir-tv2* (Fig. 1D, lower segment), offering compelling evidence of the translational potential of ORF2 within SPCs, primarily localizing to the cytosolic milieu.

Expanding our investigations, we undertook proteomic analyses, conclusively affirming the translation of both ORF1 and ORF2 in SPCs (Fig. 1E). Evidential peptide fragments were deduced from ORF1, specifically STSEQRNR and RPWSVK. In parallel, an intriguing revelation emerged regarding ORF2: the identical N-terminal sequences were identified from both *Aff3ir-tv1* and *Aff3ir-tv2*, comprising peptide fragments AESQKPVCLQETEHGR, INSRTSHNLSTDTQLASHK, MPDLLNECLVIIQPQNER, STPQAGKVNR, and VNRTRINSR. Notably, no fragments were detected emanating from the C-terminal domain of the *Aff3ir-tv1*-derived ORF2 protein. Contrarily, a pair of peptide fragments surfaced from the C-terminal region of the *Aff3ir-tv2*-derived ORF2 protein, specifically WGSVTSANRVTVR and GSRHQVLHRPHQTVTRWGSVTSANR. The representative mass spectrum of peptide fragments for AFF3IR-ORF1 and AFF3IR-ORF2 were illustrated in Fig.S2. Collectively, these empirical outcomes elucidate the translatability of both ORF1 and ORF2 within SPCs, concurrently establishing *Aff3ir-tv2* as the predominant transcript variant in this context.

We subsequently raised specific antibodies targeting LAF4IR-ORF1 and LAF4IR-ORF2, utilising peptides QRNRRPWSVKITSDC and ADHRSTPQAGKVNRC, respectively. To rigorously assess the efficacy and specificity of these antibodies, we employed mouse aortic adventitia-derived Sca1^+^ vascular progenitor cells (VPCs)^7^. Western blot (WB) analysis unveiled the presence of a band at around 7kDa by anti-ORF1 antibody, and three bands at around 16 kDa, 12 kDa and 10 kDa by anti-ORF2 antibody, observed in cultured VPCs under various conditions including exposure to laminar and disturbed flow shear stress (Fig. 1F). The pre-incubation of these antibodies with their respective peptide antigens effectively abolished the appearance of these except the 12kDa band, providing compelling evidence of the antibodies’ specificity (Fig. 1F). The 16 kDa and 10 kDa may correspond to the proteins with 151 and 109 amino acids, respectively. Indeed, ORF2 expression was highly upregulated by shear stress with laminar flow more effective (Fig.1F).

### AFF3IR-ORF1 and AFF3IR-ORF2 exhibit disparate expression patterns

To unravel the potential roles of *Aff3ir* across various biological systems, *Aff3ir*’s expression during embryonic development and in diverse adult organs was firstly investigated. As this gene was found upregulated during EC differentiation, we mainly focus on studying its role in cardiovascular system in this study. Embryonic development orchestrates the intricate maturation of the cardiovascular system at critical junctures, notably at embryonic days E8.5, E10.5, E14.5, E16.5, and E19.5. The inception of the cardiogenic plate commences around E8-E8.5, marking the genesis of cardiac myocyte cells, a pivotal milestone unfolding around E10-E10.5^15^. Subsequently, at E14-E14.5, a diverse array of vascular structures takes shape, including the aorta, vena cava, and coronary arteries, culminating in an intricate network. This intricate developmental choreography continues through the later stages of E16-E19, where both the vascular and cardiac systems undergo further refinement^16^. Our results unveiled a discernible pattern of expression for both AFF3IR-ORF1 and AFF3IR-ORF2 during embryonic development. Specifically, moderate expression levels of AFF3IR-ORF1 were noted at E8.5 and E10.5, with a significant peak observed around E14.5, followed by a decrease to near absence at E16.5. However, AFF3IR-ORF1 expression resurged markedly on the final day of embryonic development, E19.5 (Fig. 2A). Conversely, at E8.5 and E10.5, the 16 kDa AFF3IR-ORF2 exhibited a relatively low level of expression. Nonetheless, a significant upswing in AFF3IR-ORF2 expression was noted between E10.5 and E14.5 (Fig. 2A). This heightened expression level was sustained, persisting until the culmination of embryonic development at E19.5. Unlike primary VPCs cultured *in vitro*, the 10kDa band corresponding to the ORF2 protein with 109 amino acids was not detected in the tissue. Collectively, these findings illuminate a dynamic role for AFF3IR-ORF1 during the middle stages of embryonic development, notably peaking at E14.5. By contrast, AFF3IR-ORF2 appears to play a more pronounced role in the latter stages of embryonic development, underscoring the temporal intricacies of these proteins in shaping the developmental trajectory.

**Figure 2.**
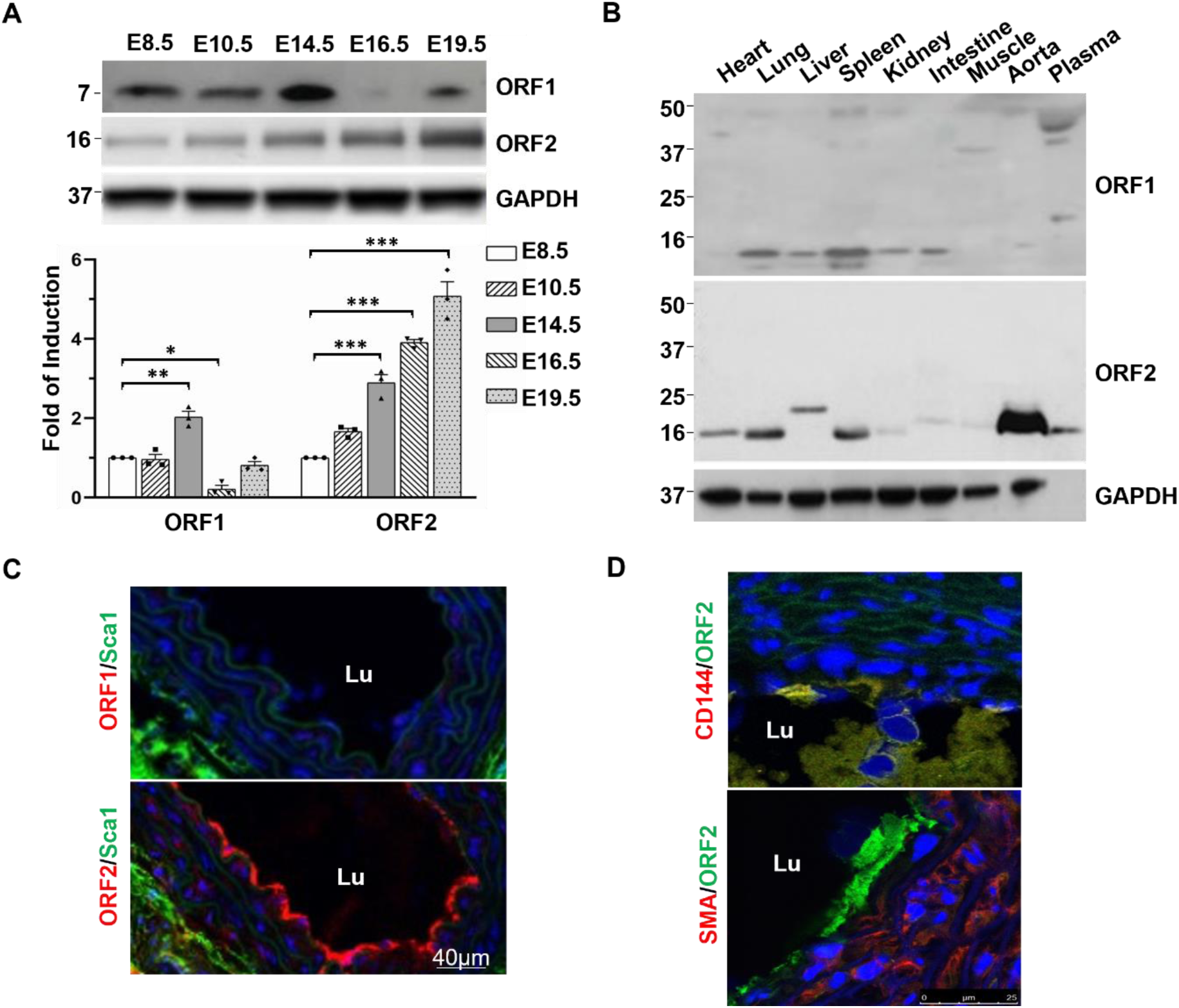
Disparate expression patterns between AFF3IR proteins. (**A**) Embryos were collected from wild type mice at stages indicated. The whole embryos were homogenized and subjected to WB. Lower panel showed the statistical analysis of the ratio of ORFs/GAPDH with that of E8.5 group set as 1.0 (n=3). (**B**) WB analysis of AFF3IR-ORFs in tissues isolated from 10-weeks old mice with GAPDH included as loading control. (**C & D**) IF staining of AFF3IR-ORFs with vascular progenitor cells marker Sca1 (**C**) or with EC marker CD144 and smooth muscle cell marker SMA (**D**) in aorta. Lu: lumen. Data presented were representative images or mean SEM using one-way ANOVA with GraphPad Prism 8 multiple comparison test. *: *p*<0.05. **: p<0.01; ***: p<0.001.

Multiple organs were collected from adult mice at 10 weeks age, followed by immunoblots. As illustrated in Figure 2B, AFF3IR-ORF1 polypeptide was discerned in select organs, including the lung, liver, spleen, and kidney, whereas it was notably low in the heart, and absent in the skeletal muscle and aorta. In stark contrast, AFF3IR-ORF2 exhibited the highest abundance within the aorta, albeit with subtle variations in size across different tissues, as depicted in Figure 2B. The predominant presence of major bands around 16 kDa strongly implies that *Aff3ir-tv2* serves as the primary isoform *in vivo*. The slight variations in band size observed among different tissues may conceivably reflect distinct posttranslational modifications impacting the AFF3IR-ORF2 protein, hinting at a nuanced regulatory layer within this intriguing molecular landscape. IF staining assays confirmed the distribution of proteins in the aorta. In line with WB result, AFF3IR-ORF1 was not detected in normal aorta (Fig. 2C). While AFF3IR-ORF2 was exclusively expressed in Sca1^+^ VPCs and ECs (Fig. 2C & 2D).

### AFF3IR-ORF2 is involved in the process of vascular injury repair

We observed the presence of only AFF3IR-ORF2 in the endothelium of the aorta. In contrast, intact femoral arteries exhibited no discernible signal for either AFF3IR-ORF1 or AFF3IR-ORF2 (Fig. 3A). However, in a femoral artery injury model^7^, where platinum wire scratching-induced injury was introduced and tissues were collected at day 3 post-surgery, a robust positive signal emerged for AFF3IR-ORF2 but not AFF3IR-ORF1 in the endothelium, lumen, and adventitia (Fig. 3A). Intriguingly, within the endothelium, AFF3IR-ORF2 localization was found external to CD31 (Fig. 3A, inset as the enlarged image), hinting at the secretion of AFF3IR-ORF2 onto the surface of endothelial cells. It is well-established that bone marrow-derived progenitor cells play a pivotal role in vascular injury repair and neointimal formation^17–19^. To explore the potential activation of AFF3IR-ORF2 during the activation of bone marrow cells, femoral bone marrow was procured from both uninjured and injured mice on day 3 after surgery. Subsequently, these cells were smeared onto glass slides and subjected to AFF3IR-ORF2 examination. While very few AFF3IR-ORF2-positive cells were observed in the bone marrow from uninjured mice, a substantial number of AFF3IR-ORF2-positive cells were detected in the bone marrow of injured mice (Fig. 3B). This observation suggests that vascular injury can activate AFF3IR-ORF2 expression in certain bone marrow cells, which might contribute to the vascular repair process following injury. Further exploration uncovered an elevation in AFF3IR-ORF2 expression within ischemic skeletal muscle tissues in a hindlimb ischemia mouse model^7^ (Fig. 3C). Collectively, these findings signify a potential association between *Aff3ir* activation and vascular remodelling, predominately mediated through the translation of the AFF3IR-ORF2 protein. Therefore, in the following studies we focused on AFF3IR-ORF2.

**Figure 3.**
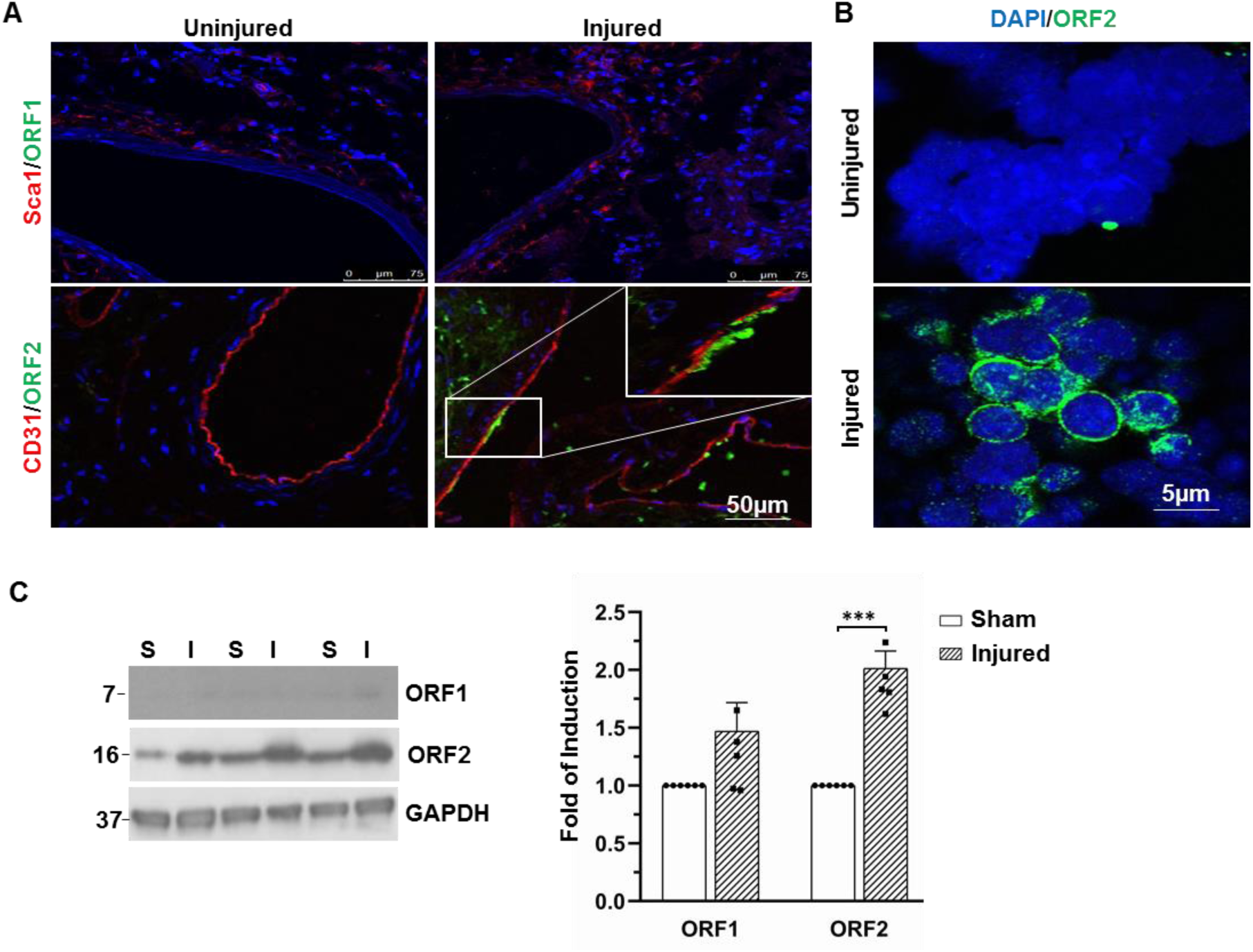
upregulation of AFF3IR-ORF2 during vascular injury repair. (**A & B**) Femoral artery injury model was introduced in wild type mice. The femoral arteries (**A**) and bone marrow tissues (**B**) were collected from uninjured or injured mice at day 3 post-surgery. The expression of relative proteins was examined by immunofluorescence staining with DAPI counterstaining nuclei. (**C**) Hindlimb ischemia model was introduced into C57BL6/J wild type mice. The skeletal muscles were collected from the uninjured or injured leg, followed by WB against relative proteins. Data presented were representative images of three independent experiments or mean SEM (n=5) using two-tail unpaired t test with GraphPad Prism 8. ***: *p*<0.001.

### AFF3IR-ORF2 promotes SPCs differentiation into ECs

As elucidated earlier, the mRNA levels of *Aff3ir* were upregulated in response to laminar flow, in parallel with the increased expression of key EC marker. Furthermore, there is a notable increase in the presence of AFF3IR-ORF2-positive cells following vascular injury. Thus, we next investigated the potential involvement of the AFF3IR-ORF2 in the context of EC differentiation through both gain-of-function and loss-of-function approaches.

As shown in Figure 4A, overexpression of AFF3IR-151aa (*Ad-Orf2*) led to a significant augmentation in laminar flow-induced mRNA levels of CD31 and CD144, both are essential EC markers. Concurrently, there was a down-regulation in the expression of smooth muscle cell marker genes, *Sm22* and calponin (*Cnn1*), within SPCs. This enhancement effect extended beyond laminar flow conditions, as evidenced by a similar observation in VEGF-induced EC differentiation from Sca1^+^ VPCs, as depicted in Figure 4B. To further study the endogenous function of AFF3IR-ORF2, we designed three siRNA fragments targeting distinct regions of the *Aff3ir* mRNA (Asi-1#: *cagagccucagucaauaca(dT)(dT)*, 335-353; Asi-2#: *gaggcaaaggaggcuacuc(dT)(dT)*, 883-901; Asi-3#: *gaccacaccagacagugac(dT)(dT)*, 1047-1065). As delineated in Figure 4C, these siRNAs exhibited varying efficacies in knocking down Aff3ir mRNA, with Asi-2# and Asi-3# displaying greater effectiveness compared to Asi-1#. Transfecting cells with Asi-2# and Asi-3# led to a notable pattern of reduction in the baseline expression of EC marker genes, specifically CD31 and CD144, within SPCs. Moreover, these siRNAs completely abrogated the laminar flow-induced upregulation of CD31 and CD144 expression (Fig. 4D), firmly implicating the active participation of AFF3IR-ORF2 in EC differentiation, especially in response to stress-induced processes.

**Figure 4.**
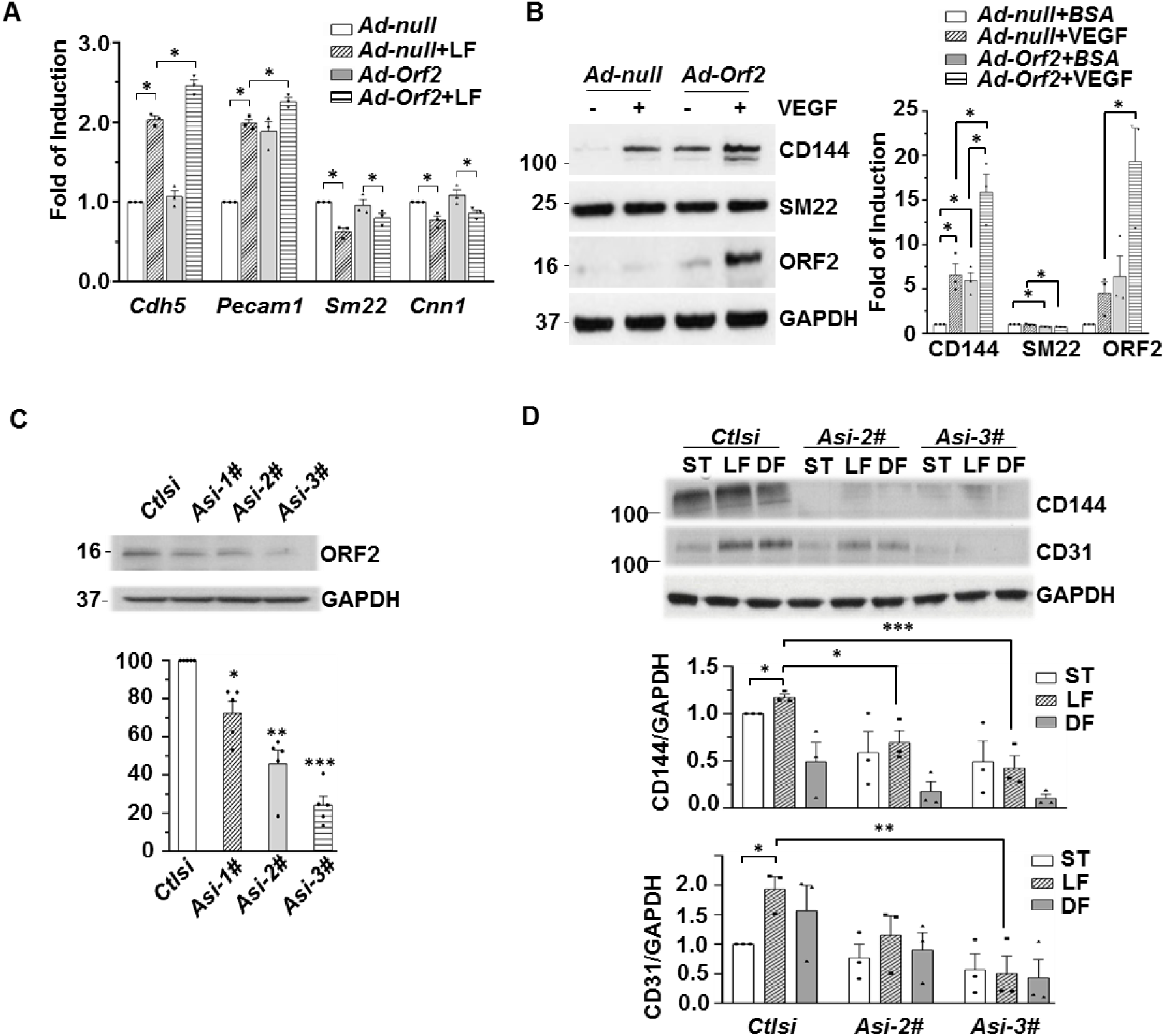
AFF3IR-ORF2 promotes SPCs differentiation into ECs. (**A**) SPCs were infected with *Ad-Orf2* at 10MOI for 24hr and then subjected to laminar flow (LF) or kept at static condition for 24hr, followed by RT-qPCR analysis of relative genes with *Gapdh* as internal control. The fold of induction was defined as the ratio of target gene to *Gapdh* comparing with *Ad-null* group. (n=3) (**B**) Sca1^+^ VPCs were infected with *Ad-Orf2* in the absence or presence of 20ng/ml VEGF for 72hr, followed by WB with antibodies indicated. *Ad-null* and BSA were included as control. The fold of induction was defined as the ratio of target protein to GAPDH with that of *Ad-null*/BSA group set as 1.0. (n=3) (**C**) SPCs were transfected with siRNAs indicated for 72hr, followed by WB analysis. Knockdown efficiency was defined as the ratio of ORF2/GAPDH with that of *ctlsi* set as 100%. (n=4) (**D**) SPCs were transfected with siRNAs indicated for 72hr and then subjected to laminar flow (LF), disturbed flow (DF) or kept at static condition (ST) for 24hr, followed by WB analysis. (n=3) Data presented were representative images or mean SEM using one-way or two-way ANOVA with GraphPad Prism 8 multiple comparison test. *: *p*<0.05. **: p<0.01; ***: p<0.001.

### AFF3IR-ORF2 inhibits SPC proliferation by sequestering MCM3 within the cytosol

In the context of cell differentiation, the inhibition of cell proliferation is critically imperative. As anticipated, overexpression of AFF3IR-ORF2 led to a pronounced suppression, while knockdown of *Aff3ir* resulted in an enhancement of proliferation in differentiated Sca1^+^ VPCs, as evaluated by cell counting (Fig. 5A) and MTT assay, respectively (Fig. 5B).

**Figure 5.**
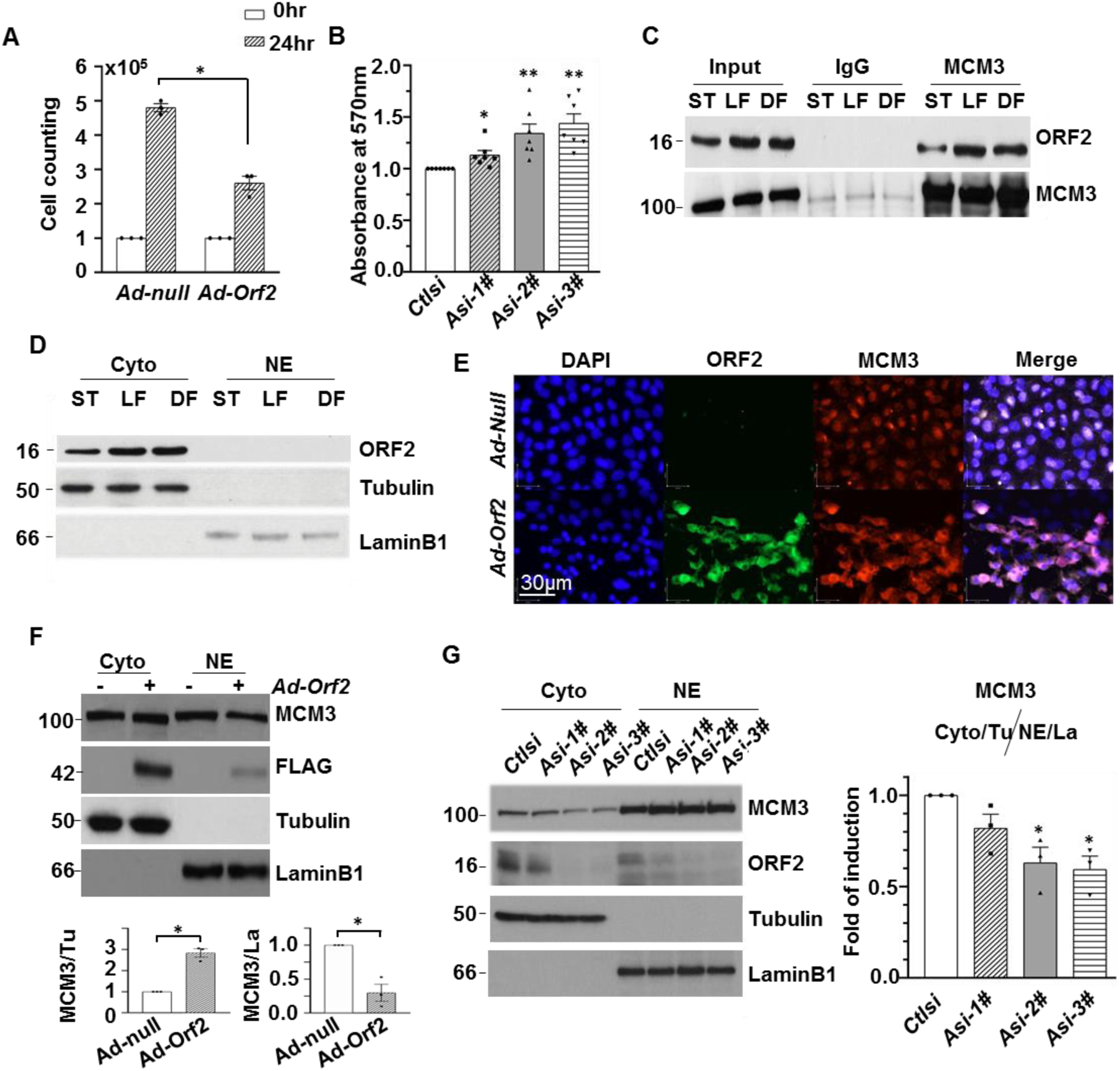
AFF3IR-ORF2 negatively regulates SPCs proliferation via retention of MCM3 in the cytosol. (**A**) Sca1^+^ VPCs were infected with *Ad-Orf2* at 10MOI for 24hr, then sub-cultured for another 24hr, followed by cell number counting (n=3). (**B**) Sca1^+^ VPCs were transfected with *Aff3ir* siRNA indicated for 48hr, and then sub-cultured for another 48hr, followed by proliferation assessment with MTT assay (n=7). (**C & D**) SPCs were subjected to laminar flow (LF), disturbed flow (DF) or kept at static condition (ST) for 24hr, followed by IP plus WB analysis (**C**), or cellular fractionation plus WB (**D**). (**E & F**) Mouse C166 cells were infected with *Ad-Orf2* at 10MOI for 24hr, followed by immunofluorescence (**E**) or cellular fractionation plus WB (**F**) (n=3). (**G**) Sca1^+^ VPCs were transfected with *Aff3ir* siRNAs followed by cellular fractionation plus WB. *Ad-null* and *Ctlsi* were included as control for the overexpression and knockdown assays, respectively. Cyto: cytosol; NE: nuclear extract. Tubulin (Tu) and Lamin B1(La) were included as loading controls for cytosolic and nuclear fractions, respectively (n=3). Data presented were representative images or mean ± SEM using two-tail unpaired t test or one-way ANOVA with Graphpad prism 8 multiple comparison test. *: *p*<0.05. **: *p*<0.01.

To explore the possible mechanisms underlying the effect of AFF3IR-ORF2 on stem cell cycle arrest, we conducted an immunoprecipitation (IP) coupled with proteomics analysis. This examination unveiled a total of 428 proteins, as illustrated in Figure S3A. Impressively, 380 of these proteins exhibited a distinct dissociation pattern in the presence of AFF3IR-ORF2 under laminar flow conditions, and further analysis was undertaken on a subset of 292 items that could be identified by the Search Tool for the Retrieval of Interacting Genes/Proteins database (STRING v11.5)^20^, revealing a network comprising 4043 edges—an observation significantly surpassing the expected 1558 (protein-protein interaction enrichment p-value < 10^-16^) (Fig S3B). Subsequent KEGG pathway analysis unveiled that a substantial portion of the dissociated proteins had functional ties to RNA processing and protein synthesis, as depicted in Figure S3C. This revelation strongly suggests a pivotal role for AFF3IR-ORF2 in post-transcriptional regulation and its regulatory influence in orchestrating stem cell cycle arrest.

Among the proteins uniquely identified in the laminar flow-treated samples, one of particular significance was the minichromosomal maintenance complex component 3 (MCM3), a highly conserved protein known for its pivotal role in promoting the initiation of DNA replication^21^. IP plus WB analysis showed the physical interaction between MCM3 and AFF3IR-ORF2 in SPCs, which was further potentiated under laminar flow conditions (Fig. 5C). Moreover, cellular fraction analysis unveiled that the AFF3IR-ORF2 bands were predominantly abundant within the cytosol fraction (Fig. 5D). Immunofluorescence (IF) staining further substantiated these findings, revealing a striking co-localization of AFF3IR-ORF2 and MCM3 within the cytosol in ECs (Fig. 5E). This pattern persisted upon the overexpression of AFF3IR-ORF2 using *Ad-Orf2* in mouse C166 EC cell line (Fig. 5F), by contrast, knockdown of *Aff3ir* resulted in a diminished retention of MCM3 within the cytosol in Sca1^+^ VPCs (Fig. 5G). Notably, it’s worth mentioning that the FLAG-tagged ORF2 protein predominantly appeared as a 42 kDa band in C166 cells, unlike the 16 kDa observed in stem cells. This discrepancy suggests potential cell type-specific post-translational modifications impacting ORF2.

Building upon our previous studies, which elucidated that laminar flow could upregulate p21^waf1^ via histone deacetylase-mediated p53 deacetylation, resulting in cell cycle arrest and differentiation^9, 22^, we discovered that the overexpression of AFF3IR-ORF2 led to an increase in the expression of p53 and p21^waf1^ (Fig. S4). These collective findings strongly indicate that AFF3IR-ORF2 exerts a negative modulatory effect on cell proliferation through the retention of MCM3 within the cytosol and the upregulation of the p53/p21^waf1^ pathway. Consequently, this regulatory milieu enhances the capacity of stem cells to embark on the journey of differentiation towards ECs.

## Discussion

This report unveils a novel nested protein-encoding duocistronic gene, *Aff3ir*, situated within the opposite strand of intron 6 of the mouse transcription factor gene *Aff3*. *Aff3ir* encodes a 45-amino acid sequence via Orf1 (AFF3IR-ORF1), and a 109-amino acid or 151-amino acid sequence via Orf2 within *Aff3ir-tv1* or *Aff3ir-tv2* (AFF3IR-ORF2). Significantly, AFF3IR-ORF1 and AFF3IR-ORF2 exhibit disparate expression patterns and distinct distribution *in vivo*. AFF3IR-ORF2 with 151-amino acid sequence plays a critical role in EC differentiation by orchestrating a shift in the status of VPCs, redirecting them from proliferation towards differentiation into ECs.

The cDNA fragment of *Aff3ir* is firstly reported as a predicted non-coding gene by murine RIKEN integrated sequence analysis^23, 24^. As a special type of overlapping gene that is entirely embedded within the chromosomal region occupied by another gene^25^, *Aff3ir* lies within a large intron of *Aff3*, which is a common form for protein-coding nested gene in eukaryotes^26^. The discovery of the *Aff3ir* gene represents a valuable model system to illuminate the presence of other yet-to-be-identified genes and their intricate transcriptional regulations within vast intronic regions. Utilizing opposite DNA strands for their transcriptions, *Aff3* and *Aff3ir* may employ either an intron-bypass transcription mechanism or a temporary transcription mechanism. In the former scenario, transcription of *Aff3* circumvents the vast introns, including intron 6, thus allowing simultaneous yet independent transcription of both *Aff3* and *Aff3ir*. This mode has the added benefit of conserving energy during the transcription of genes replete with large introns or those that necessitate skipping multiple exons^27^. In the latter mode, transcription of *Aff3* and *Aff3ir* unfolds at distinct time points, avoiding the formation of double-stranded RNAs between *Aff3ir* and *Aff3* precursor that contains intron 6. This mechanism serves to prevent the activation of the cellular autonomous immune system, thereby safeguarding both RNA molecules from destruction^28^. Elucidating the intricate mechanisms governing the transcription of *Aff3* and *Aff3ir* will increase our understanding of transcription regulation, particularly in the context of genes characterized by extensive intronic sequences.

The presence of multiple transcript variants in multiexon genes, arising from alternative splicing in cell lines^29^, leads to the detection of both proteins with 109-amino acid and 151-amino acid sequences in SPCs. However, in primary tissues, a usual scenario involves only one dominant transcript per gene^30^, resulting in the detection of a single band of AFF3IR-ORF2 in spite of the fact that the size of band varies among different organs, suggesting the possibility of additional transcript variants.

Lining the inner surface of blood vessels, vascular endothelium plays a pivotal role in regulating vessel tone and permeability, while dysfunction of ECs can lead to issues such as vessel dilation and haemorrhage^31–33^. Vascular remodelling, on the other hand, is crucially orchestrated by SPCs, which can originate from local resident progenitors or mesenchymal stem cells sourced from the bone marrow^34–36^. We found that, in response to vessel injury, the upregulation of AFF3IR-ORF2 within the vasculature and bone marrow may contribute to vascular remodelling. Furthermore, it is reasonable to consider AFF3IR-ORF2 as a potential marker for endothelial progenitor cells, a notion supported by the positive correlation between its expression levels and the EC differentiation process.

Cell cycle arrest is a pivotal prerequisite for stem cell differentiation, an intricately orchestrated process which the underlying mechanisms are complex and still not fully understood. Our findings indicate that AFF3IR-ORF2 has an inhibitory role in modulating SPC proliferation through its interaction with MCM3. MCM3 belongs to the highly conserved family of mini-chromatin maintenance proteins^21^. This family is renowned for its role as DNA replication licensing factors, crucial for binding to DNA replication origins and orchestrating the unwinding of densely packed chromatin structures. Elevated levels of nuclear MCM3 are often linked to heightened cancer cell proliferation^21, 37^. Our data demonstrate has unveiled a compelling interaction between AFF3IR-ORF2 protein and MCM3 in the cytosol, particularly under the influence of laminar flow stress. This interaction leads to the retention of MCM3 in the cytosol, subsequently the decreased abundance of nuclear MCM3. As a result, this in turn hinders DNA replication and instigates cell cycle arrest.

It is worth noting that while AFF3IR-ORF2 is predominantly located in the cytosol, trace amounts are also discernible in the nucleus. This prompts further exploration into the possibility of AFF3IR-ORF2 engaging with MCM3 within the nucleus. Beyond its role in DNA replication, MCM3 may function as a co-activator or co-repressor, modulating gene transcription. Evidence supporting this notion includes findings that MCM proteins, including MCM3, can suppress hypoxia-inducible factor 1 activity through direct interaction^38^. Furthermore, MCM3 has been implicated in hematopoietic stem cell differentiation differentiation^39^. AFF3IR-ORF2 is an alkaline protein, with 51 alkaline residues (Arg, His and Lys) versus 19 acidic residues (Asp and Glu). The net positive charge enables AFF3IR-ORF2 bind to DNA or RNA molecules. This broader perspective underscores the potential impact of the association between AFF3IR-ORF2 and MCM3 on gene transcription regulation.

In conclusion, our discovery reveals the novel discovered *Aff3ir* gene as a regulator in stem cell differentiation, operating mainly through spatial interactions with MCM3. The complicated mechanisms governing this gene undoubtedly hold the potential to unlock new frontiers in our understanding. These include the exploration of hitherto undiscovered genes nestled within extensive introns, the quest for the human analogue of AFF3IR, the unravelling of transcriptional regulations within these expansive genetic domains, and the elucidation of their roles in fundamental physiological and pathophysiological processes, notably including the intricate domain of vascular remodelling.

## Materials and Methods

### Materials

All cell culture media and sera were purchased from Thermo Fisher Scientific (Waltham, MA, USA), whereas cell culture supplements and growth factors were purchased from Sigma (St. Louis, MO, USA). The antibodies against AFF3IR-ORF1 in mouse and against AFF3IR-ORF2 in rabbit were synthesized by GenScript (Piscataway, NJ, USA). The antibodies against Sca-1 (ab51317), CD144 (ab205336), SM22 (ab14106), p53 (ab26), Caspase-3 (ab4051) and Lamin B1 (ab16048) were purchased from Abcam (Cambridge, UK). FLAG (F1804), p21^waf^^1^ (MABE325) and tubulin (T6199) were purchased from Sigma (Dorset, UK). CD31 (af3628) was purchased from R&D systems (Abingdon, UK), GAPDH (sc-25778) and MCM3(sc-390480) were purchased from Santa Cruz Biotechnology (Dallas, TX, USA). All secondary antibodies and ECL reagents were purchased from Dakocytomation (Glostrup, Denmark). Routine chemicals were purchased from Sigma.

### Cell culture

Mouse ES cells (ESCs, ES-D3 cell line, CRL-1934) were purchased from ATCC and maintained in stem cell culture medium [DMEM (ATCC, Rockville, MD, USA) supplemented with 10ng/mL recombinant human leukaemia inhibitory factor (LIF; Chemicon, Temecula, CA, USA), 10% foetal bovine serum (FBS, ATCC), 0.1mmol/L 2-mercaptoethanol(2ME), 100U/mL penicillin, and 100µg/mL streptomycin] as describe previously^40^. Mouse aorta adventitia-derived Sca1^+^-vascular progenitor cells (VPCs) were isolated and cultured as described previously ^7^. The Sca-1^+^-VPCs were maintained in stem cell culture medium and passaged every two days at a ratio of 1:4 by Trypsin-EDTA (0.25%) phenol red (ThermoFisher, UK). Up to 5 passages were used in this study. HEK293 (for adenovirus amplification) and mouse C166 cells (endothelial cell line, CRL-2581) were purchased from ATCC. HEK293 and C166 cells were maintained in DMEM (high glucose, Gibco) supplemented with 10% FBS (Giboco), L-glutamine (2mM, Giboco), penicillin (100U/mL) and streptomycin (100µg/mL).

### Cell differentiation

Mouse ESCs were seeded on 5µg/mL Fibronectin (FN)-coated glass slide or dishes (Ø10cm) or 25mL-flasks in differentiation medium [DM, αMEM supplemented with 10% FBS, 0.05mmol/L 2ME, penicillin (100U/mL) and streptomycin (100µg/mL)] for 3 days with medium refreshment every other day to let the cells spontaneously differentiate to obtain stem/progenitor cells (SPCs). For AFF3IR-ORF2 overexpression, the 3-day differentiated ESCs derived SPCs were infected with *Ad-null* or *Ad-Orf2* virus at 10 multiplicity of infection (MOI) for 24hr. The cells on slides were subjected to laminar shear at 12dynes/cm^2^ in a rectangular flow system for 24hr as described previously^9^, followed by RNA extraction and quantitative reverse transcriptase-polymerase chain reaction (qRT-PCR) or microarray analysis by Dr Shu Chien’s group (University of California, San Diego, CA USA). The cells in dishes were subjected to rotation at 150rpm for 24hr by a PSU-10i orbital shaker (Grant-Bio, UK), followed by protein analysis. The outer 2cm area was defined as laminar flow (LF), while the remaining central part was defined as disturbed flow (DF). The SPCs derived from 3-day differentiated ESCs in flasks were infected with adenovirus at 10 MOI or transfected with siRNA for 24hr, and then continuously cultured in DM supplemented 2% FBS and 20ng/mL mouse VEGF-165 for 7 days with medium refreshment every other day, followed by WB analysis.

### RNA extraction and quantitative RT-PCR

Total RNAs were extracted using RNeasy Mini kit (Qiagen, Manchester, UK) according to the protocol provided. One μg of RNA with its genomic DNA ablated was converted into cDNA as instructed by the QIAGEN OneStep RT-PCR Kit (Qiagen, UK). Twenty ng cDNA (relative to RNA amount) was subjected to quantitative PCR with a SYBR green dye–based PCR amplification and detection master mix (ThermoFisher Scientific) in Eppendorf MarsterCycler gradient S machine (Eppendorf, Enfield, CT, USA). The primers were designed using *ncbi primer pickup* software and synthesized by Thermofisher (UK). The primer sequences include: *Gapdh* (5’-cga ctt caa cag caa ctc cca ctc ttc-3’ vs 5’-tgg gtg gtc cag ggt ttc tta ctc ctt-3’), *Aff3ir* (5’-ggt ctg gag ttg cct tca-3’ vs 5’-tgg aac ctc tcc tct gtc tt-3’), *Aff3* (5’-cta cca caa cc acta cca cta c-3’ vs 5’-cca ggt gac tgc tat cca taa g-3’), *Cah5* (5’-gtg cct gaa gac atc cga gtg-3’ vs 5’-gac ctc tgt cac tgg tct tgc-3’), *Pecam1* (5’-gga gtg cct tgt gga cat cag-3’ vs 5’-tgc acg gtg acg tat tca ctc-3’) and *β-actin*(5’-cac acc tgg gac gac atg gag-3’ vs 5’-ttc atg agg tag tga gtc tgg-3’).The fold of induction was defined as the ratio of the difference between target gene and *Gapdh* or *β-actin* (internal control) with that of *Ad-null*/static set as control at 1.0.

### Plasmid construction

The full length *Aff3ir* mRNA was cloned using commercially available 5’-RACE and 3’-RACE kits (ThermoFisher Scientific) according to the protocols provided. The FLAG tag sequence was inserted downstream the ATG start codon of *Orf2* of *Aff3ir-tv2* by high fidelity PCR. The full sequence was then cloned to pShuttle 2 vector to obtain *pShuttle2-FLAG-Aff3ir-tv2* plasmid.

The cDNA sequence for the 151-aa of ORF2 were synthesized (Genscript) with FLAG tag sequence inserted downstream the ATG start codon and cloned to pShuttle 2 vector, designated as pShuttle2-*Orf2*. Accordingly, *Ad-Orf2* was created using adenoviral X system (Clontech, Mountview, CA, USA) and viral particles were created and amplified in HEK293 cells according to the protocol provided. *Ad-null* was generated with pShuttle2 vector.

### *siRNA* knockdown

The control siRNA (*5′-ccucaucgccuggcauugatt-3′*) and the siRNAs for *Aff3ir* (*Asi-1#:* 5’-*cagagccucagucaauacatt-3’*, 335-353; *Asi-2#:* 5’-*gaggcaaaggaggcuacuctt-3’*, 883-901; *Asi-3#:* 5’-*gaccacaccagacagugactt*, 1047-1065) were purchased or synthesized from Ambion as double stand RNAs. The transfection was performed using the GenMute Reagent (SignaGen Laboratories, Frederic, USA) with the provided protocol modified. Briefly, 150pmol siRNA (3µL of 50µmol/L) was diluted in 1× buffer and mixed with 12 µL and incubated at room temperature for 15min. During this period, the cells were detached by trypsin, and single cell suspension at 5×10^5^cells/mL was prepared with the differentiation medium. The siRNA mixture was then mixed with 1ml cell suspension and incubated on a rotator at 50rpm at 37°C for 5 hrs to let transfection occur. The transfected cells were then seeded in FN-coated flasks in differentiation medium for three days with medium refreshment every other day, followed by differentiation or proliferation assays accordingly.

### Western blot, immunoprecipitation and immunofluorescence staining

WB, IP and IF were performed according to standard procedures described elsewhere. For IP, 1mg cell lysate or 500 µg cytosolic or nuclear fraction were incubated with 2μg antibody or normal IgG and pulled down by 5µL protein G-magnetic beads (Bio-Rad). Samples were then pulled down by protein-G-magnetic beads and subjected to WB or proteomics analysis. For the peptide blocking assays, the antibody was pre-incubated with blocking peptide at a ratio of 1:1 (1μg peptide versus 1μg antibody) at 4 °C overnight. To reduce background, the incubation of primary antibodies was performed at 4 °C overnight for both procedures. Primary antibodies were used at a concentration of 0.5-1 µg/mL for WB and 2-10 µg/mL for IF. Secondary antibodies were used at 1:3000 dilution for WB, 1:2000 dilution of IF. In the peptide blocking assays in IF, secondary antibody was used at 1:500 dilution.

### Proteomics analysis

For small molecular proteins, proteins isolated from 3 day-differentiated ESCs were applied to 4-12% SDS-PAGE gel, followed by gel cutting between the indicative dye band and 37kDa band. For immunoprecipitated samples, the eluted samples were applied to 4-12% SDS-PAGE gel and run until the indicative dye band migrated around 1.5cm, followed by gel cutting between the well and the indicative dye band. The cut gets were subjected to in-gel trypsin digestion and tandem liquid chromatography mass spectrometry.

### Cell proliferation, cell survival and MTT assay

Sca1^+^-VPCs were infected with *Ad-Orf2* (*Ad-null* included as control) at 10MOI for 24hr, followed by cell counting using a multisizer 3 coulter counter (Beckman Coulter), according to the manufacturer’s instructions^41^. SPCs or Sca1^+^-VPCs were transfected with *Aff3ir siRNA* as described above for 48hr. The cells were then seeded in 24 well plates at 2×10^4^cells/well/0.5ml medium and cultured for 48hr. The cells were then subjected to proliferation assay using CellTiter 96® Non-Radioactive Cell Proliferation Assay (MTT) (Promega, Madison, USA) with protocol provided.

### Animal study

All animal experiments were performed according to the protocols approved by the Institutional Committee for Use and Care of Laboratory Animals under the UK Home Office Project License PPL*70/7266* (femoral artery injury model and hind limb ischemia model).

Wild type C57bl/6 (8-10 weeks) were purchased from Charles River (West Sussex, UK) and maintained in King’s College London animal facility according to standard procedures. The mice were used for embryonic and adult tissue collection, Sca1^+^-VPCs cell isolation^7^, femoral artery injury and hindlimb ischemia surgery under 2% isoflurane anaesthesia as described previously ^41^. The injured vessels and bone marrow were harvested at day 3 post-surgery for femoral artery injury model. The adductor muscle tissues were collected at day 7 post-surgery from the hindlimb ischemia model. Six mice were used for each group and uninjured mice were included as sham control. Cryo-sectioned IF or WB was performed on these tissues.

### Statistical Analysis

Data were characterised as mean ± standard error of the mean (SEM). All data were analysed using GraphPad Prism 8 software with t-test, one-way or two-way analysis of variance (ANOVA) followed by Dunnett’s multiple comparison tests. The significance was depicted by asterisks, *: p<0.05, **: p<0.01, ***: p<0.001, ns: no significant difference. A value of p < 0.05 was considered significant.

## Source of funding

This study was supported by funding from British Heart Foundation project grants PG12/11/29408 (LZ) and PG/22/11055 (MZ), PhD Studentship FS15/74/31669 (LZ) and YZ is partially supported by China Scholarship Council.

## Author contribution

Y. Z., M. E. and Y. L. contributed to experimental design, performance, data analysis and paper writing. X. B., L. H., A. A., X. Y., S. L. and Y. T. contributed to experimental performance. Y. G., and A. M. contributed to data analysis; T. C. and A.M.S. contributed to manuscript writing. T. L., M.Z. and L. Z. contributed to experimental design, data analysis and manuscript writing. All authors were involved in critical evaluation and intellectual contribution to the manuscript.

## Disclosures

The authors declare no competing interests.

## Notes

### Competing Interest Statement

The authors have declared no competing interest.

