## Supplemental Figures for "A novel nested gene *Aff3ir* participates in vascular remodelling by enhancing endothelial cell differentiation in mice"

**A** **transcript variant 1, 2059nt**

accccttcccttaacaccaaaacagctgaccctgagatggaggcttacatttccactcccagctgaccatcacctctgcgctaaggccaaaacacagcatgtactg
gggagaagcaagcaggcaagactggagaagcacttcagaacaaagaaacagacgtccttggagcgtcaagatcaccagtgacaggaaacggattctaga
aagcagatactggaactctgctgcagtgtaaaaaaaacaataggagatgagcaagcagctgataacgactcgggcgtgtgatccagttacaaaagcaagccg
ccaccgccacccccaccccggctaactgcagagcctcagtcaatacatccaggaaagaccacagaatcaatgacaaggctaatttaacagaattaagaagtg
gcgttagggaaaaaaaaattaaggaagtccacataactgcaattaccatatccttttaatgttgcttagcaacgatatataaaaatgagatataatgtcgggaactta
gtttcccggacagaaacagtgcctgcctcagcagggctgtaactcacttttctccaactgcaactccgtgggttgctgtgggtggtgctacttctgtaatactttctcttac
acggtcatgtgtttgtgttagtgcattaaaatgccggatttgcttaatgagtgcctggtgataatacagccacagaatgagagggctcttattccatttgcttctgcagatc
acaggtccactccacaggcaggaaaagtcaacagaaccagaataaactcaaggacatcacataatctatccacagacactcagctagcaagccataaagct
gaaagtcaaaagcctgtctgtcttcaagaaactgaacacgggaggcaaaggaggctactcaagattgttgctgttgtgtgtgggagtggtctggagttgccttcattt
aagtttacaaggcactttgaatatccatctttcttgtgagaacctcagacctccattgcaagacagaggagaggttccaggcatcaggttctccacagaccacacca
gacagtgacaagatgggggagtgttacttctgcaaacagagtcactgtgcggctgagaacaccagaagatgtggtaaagagacatcctggctgttgctctccag
agtctttgccatcctaaaggaggggtgctatgtggtggcccagcctcctgtgaacgatgggtttcctactagacaggaggaggaggtcaaccccaagggaagcc
atgccacagtgattcctgaaatgggaccaatcccggttggtgaccagccctgaaggtgagacacctgcagaagctgaggagtgtggcacacctgctggcccttc
ctgatagagccacatgggactcagcctggagcagcctctggctacagcatcctttcaatgtaagacctgctaagccccaggatccccagatgctcagtgtacgca
tccttcatgactctcactggcaaaagatatggaaagagacagtcactgatggatggaaagggactgtgtgagtgtaaacacgtaacgggaagtcctgcgcctctc
ttaattgcccgacacaattcccactacgtgcataaactgcagttcccattaacagcaccatgaataataatcagtttactttctgactcgtatctaaatataccttctatat
gatttcatcttttatgcagctgtttagaagcagaacatctaattacagttcctattagtcaacattaatttttttccctttgccacggagcatatgtaatgactgattttctaaa
aacagaatttttttagcatatggaacaaattgtgttattattaatgcattcccagatgactctgcatttcaattcccttaattatttagtcattgctgattatgtcttttaacattc
attatcccatctcttgaaacacaggcagtttattcattaagtgtcaaaatggaaagcttgacctccaagaaggaaaaaaaaaaagataatcaggaaaaccgaaaa
aaaaaaaaaaaaaaaaaaaaaaaaaaaaa

**ORF1: 98~235** MYWGEASRQDWRSTSEQRNRRPWSVKITSDRKRILESRYWNSAAV

**ORF2: 659~988** MPDLLNECLVIIQPQNERALIPFASADHRSTPQAGKVNRTRINSRTSHNLSTDTQLASHKAES
QKPVCLQETEHGRQRRLLKIVAVVCGSGLELPSFKFTRHFEYPSFL

**B** **transcript variant 2, 1996nt**

accccttcccttaacaccaaaacagctgaccctgagatggaggcttacatttccactcccagctgaccatcacctctgcgctaaggccaaaacacagcatgtactg
gggagaagcaagcaggcaagactggagaagcacttcagaacaaagaaacagacgtccttggagcgtcaagatcaccagtgacaggaaacggattctaga
aagcagatactggaactctgctgcagtgtaaaaaaaacaataggagatgagcaagcagctgataacgactcgggcgtgtgatccagttacaaaagcaagccg
ccaccgccacccccaccccggctaactgcagagcctcagtcaatacatccaggaaagaccacagaatcaatgacaaggctaatttaacagaattaagaagtg
gcgttagggaaaaaaaaattaaggaagtccacataactgcaattaccatatccttttaatgttgcttagcaacgatatataaaaatgagatataatgtcgggaactta
gtttcccggacagaaacagtgcctgcctcagcagggctgtaactcacttttctccaactgcaactccgtgggttgctgtgggtggtgctacttctgtaatactttctcttac
acggtcatgtgtttgtgttagtgcattaaaatgccggatttgcttaatgagtgcctggtgataatacagccacagaatgagagggctcttattccatttgcttctgcagatc
acaggtccactccacaggcaggaaaagtcaacagaaccagaataaactcaaggacatcacataatctatccacagacactcagctagcaagccataaagct
gaaagtcaaaagcctgtctgtcttcaagaaactgaacacgggaggcaaaggaggctactcaagattgttgctgttgtgtgtgggagtggtctggagttgccttcattt
aagaggagaggttccaggcatcaggttctccacagaccacaccagacagtgacaagatgggggagtgttacttctgcaaacagagtcactgtgcggctgagaa
caccagaagatgtggtaaagagacatcctggctgttgctctccagagtctttgccatcctaaaggaggggtgctatgtggtggcccagcctcctgtgaacgatgggt
ttcctactagacaggaggaggaggtcaaccccaagggaagccatgccacagtgattcctgaaatgggaccaatcccggttggtgaccagccctgaaggtgag
acacctgcagaagctgaggagtgtggcacacctgctggcccttcctgatagagccacatgggactcagcctggagcagcctctggctacagcatcctttcaatgt
aagacctgctaagccccaggatccccagatgctcagtgtacgcatccttcatgactctcactggcaaaagatatggaaagagacagtcactgatggatggaaag
ggactgtgtgagtgtaaacacgtaacgggaagtcctgcgcctctcttaattgcccgacacaattcccactacgtgcataaactgcagttcccattaacagcaccatg
aataataatcagtttactttctgactcgtatctaaatataccttctatatgatttcatcttttatgcagctgtttagaagcagaacatctaattacagttcctattagtcaacatt
aatttttttccctttgccacggagcatatgtaatgactgattttctaaaaacagaatttttttagcatatggaacaaattgtgttattattaatgcattcccagatgactctgc
atttcaattcccttaattatttagtcattgctgattatgtcttttaacattcattatcccatctcttgaaacacaggcagtttattcattaagtgtcaaaatggaaagcttgacct
ccaagaaggaaaaaaaaaaagataatcaggaaaaccgaaaaaaaaaaaaaaaaaaaaaaaaaaaaaa

**ORF1: 98~235** MYWGEASRQDWRSTSEQRNRRPWSVKITSDRKRILESRYWNSAAV

**ORF2: 659~1114** MPDLLNECLVIIQPQNERALIPFASADHRSTPQAGKVNRTRINSRTSHNLSTDTQLASHKAES
QKPVCLQETEHGRQRRLLKIVAVVCGSGLELPSFKRRGSRHQVLHRPHQTVTRWGSVTSANRVTVRLRTPED
VVKRHPGCCSPESLPS

**Fig. S1: The nucleotide and deduced amino acid sequences for *Aff3ir* transcript variants.** The orf1 and orf2 nucleotide sequences were underlined. The identical amino acids sequences of ORF2 protein from the two transcript variants were underlined as well.

### AFF3IR-ORF1 peptide-P9

Sequence: **STSEQRNR**, Charge: +2, Monoisotopic m/z: 489.24000 Da (-1.59 mmu/-3.25 ppm), MH+: 977.47272 Da, RT: 71.4742 min,
Identified with: Sequest HT (v1.17); XCorr:1.53, Percolator q-Value:0.0e0, Percolator PEP:1.8e-1,
Fragment match tolerance used for search: 0.6 Da
Fragments used for search: -$H_2O$; y; -$NH_3$; y; b; b; -$H_2O$; b; -$NH_3$; y
Proteins (1): - P00005 peptide_mouse

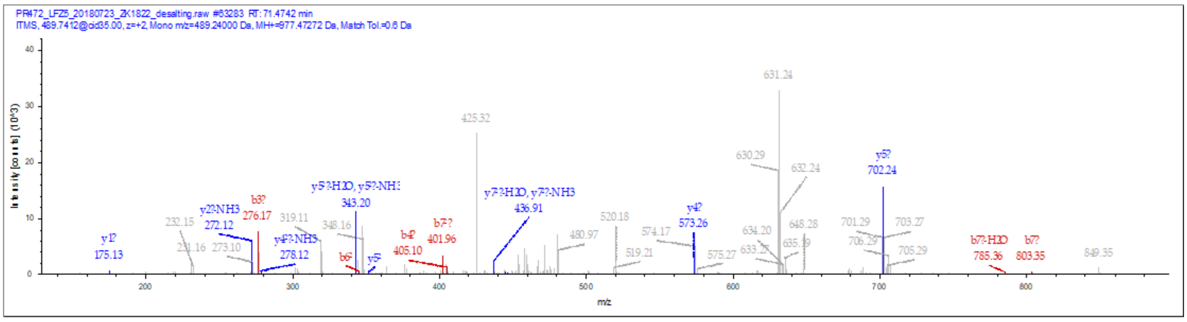

### AFF3IR-ORF2 peptide-P1

Sequence: ***AESQKPVCLQETEHGR***, C8-Carbamidomethyl (57.02146 Da)
Charge: +2, Monoisotopic m/z: 934.95343 Da (+3.83 mmu/+4.09 ppm), MH+: 1868.89958 Da, RT: 28.6622 min, Identified with: Sequest HT (v1.17); XCorr:2.58,
Fragment match tolerance used for search: 0.6 Da

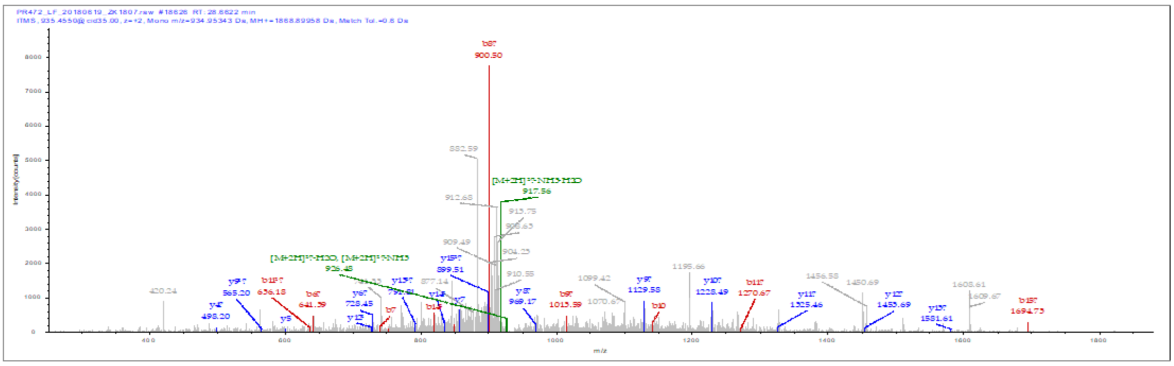

### AFF3IR-ORF2 peptide-P7

Sequence: ***WGSVTSANRVTVR***, Charge: +2, Monoisotopic m/z: 716.88232 Da (-4.08 mmu/-5.7 ppm), MH+: 1432.75737 Da, RT: 34.0684 min, Identified with: Sequest HT (v1.17); XCorr:2.09,
Fragment match tolerance used for search: 0.6 Da,

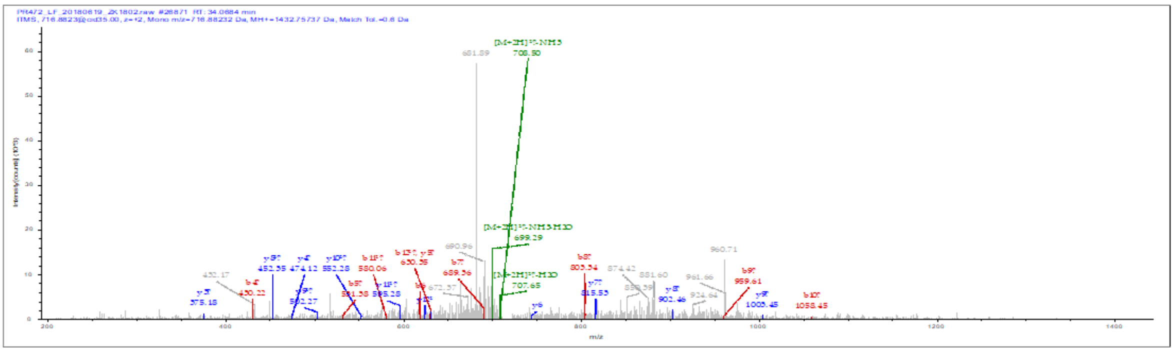

**Fig.S2: Representative mass spectrum of peptide fragments identified by proteomics analysis.** P# corresponds to that in Fig. 1E.

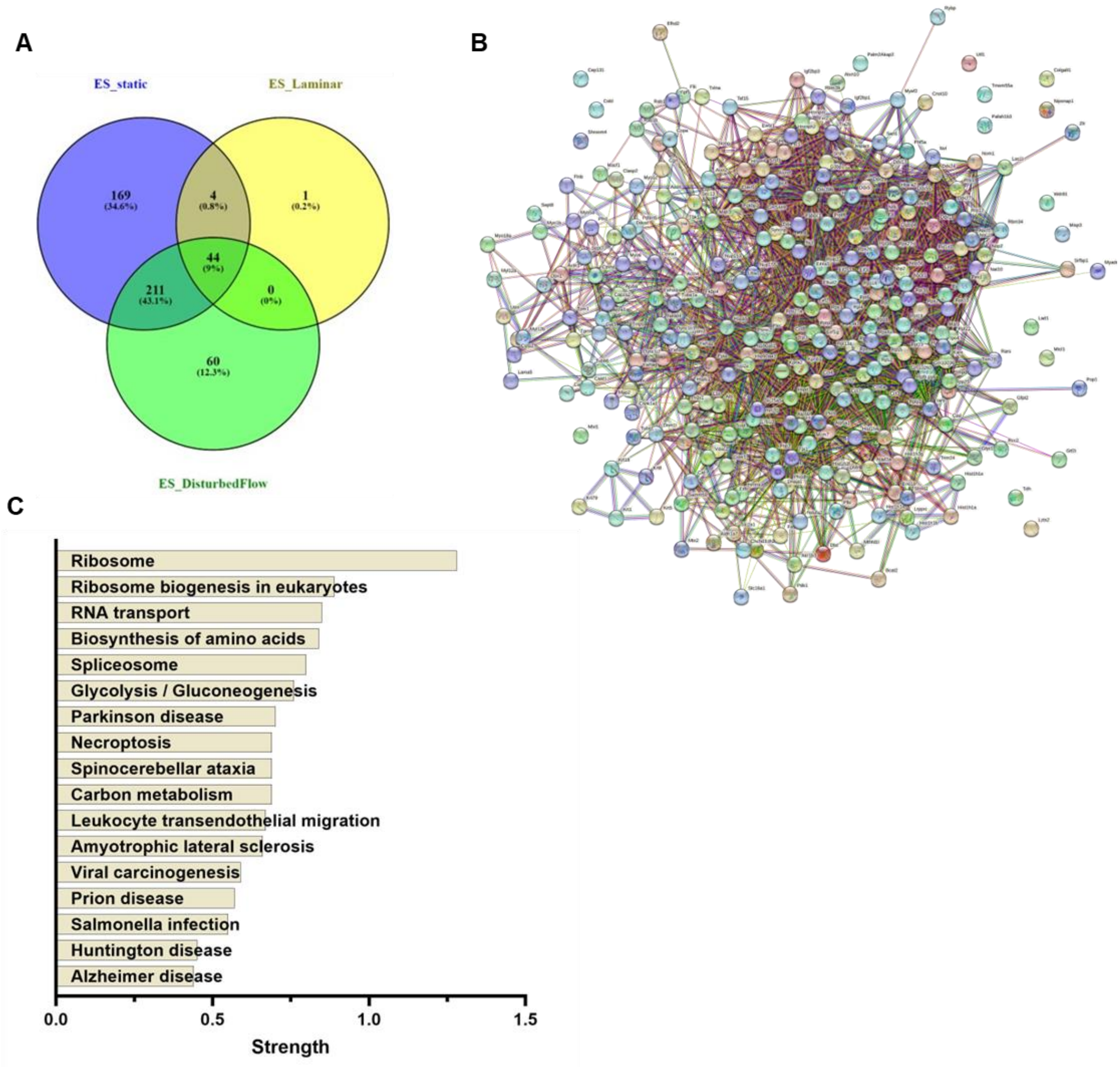


**Fig.S3: AFF3IR-ORF2 associated proteins by IP with anti-AFF3IR-ORF2 plus proteomics analysis**. (**A**) Venn diagram of associated proteins. (**B**) protein-protein interaction analysis by the Search Tool for the Retrieval of Interacting Genes/Proteins database (STRING v11.5) (**C**) KEGG pathway analysis of 292 identified associated proteins in STRING database.

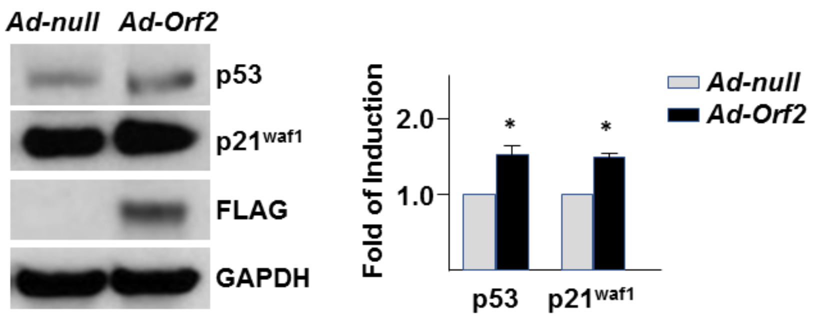


**Fig.S4: p53/p21waf1 pathway may be involved in AFF3IR-ORF2-mediated cell proliferation suppression.** SPCs were infected with *Ad-Orf2*, followed by WB with antibodies indicated. *Ad-null* was included as control. The fold of induction was defined as the ratio of target protein to GAPDH with that of *Ad-null* group set as 1.0. (n=3) Data presented were representative images or mean SEM using two-tail unpaired t test with GraphPad Prism 8 multiple comparison test. *: $p$<0.05.
